# CHIPS: A Snakemake pipeline for quality control and reproducible processing of chromatin profiling data

**DOI:** 10.1101/2021.03.09.434676

**Authors:** Len Taing, Clara Cousins, Gali Bai, Paloma Cejas, Xintao Qiu, Zach Herbert, Myles Brown, Clifford A. Meyer, X. Shirley Liu, Henry W. Long, Ming Tang

## Abstract

**Motivation:** The chromatin profile measured by ATAC-seq, ChIP-seq, or DNase-seq experiments can identify genomic regions critical in regulating gene expression and provide insights on biological processes such as diseases and development. However, quality control and processing chromatin profiling data involve many steps, and different bioinformatics tools are used at each step. It can be challenging to manage the analysis.

**Results:** We developed a Snakemake pipeline called CHIPS (<u>CH</u>romatin enr<u>i</u>chment <u>P</u>roce<u>s</u>sor) to streamline the processing of ChIP-seq, ATAC-seq, and DNase-seq data. The pipeline supports single- and paired-end data and is flexible to start with FASTQ or BAM files. It includes basic steps such as read trimming, mapping, and peak calling. In addition, it calculates quality control metrics such as contamination profiles, PCR bottleneck coefficient, the fraction of reads in peaks, percentage of peaks overlapping with the union of public DNaseI hypersensitivity sites, and conservation profile of the peaks. For downstream analysis, it carries out peak annotations, motif finding, and regulatory potential calculation for all genes. The pipeline ensures that the processing is robust and reproducible.

**Availability:** CHIPS is available at https://github.com/liulab-dfci/CHIPS

## 1 Introduction

Protein-DNA binding interactions are fundamental to gene regulation and involved in regulating disease processes. However, the methods of investigating these interactions through ATAC-seq, ChIP-seq, and DNase-seq experiments generate data that require extensive processing before biological interpretation (Furey, 2012). Chromatin profiling using sequencing technology can also generate bias, which needs to be mitigated before interpreting the biological significance (Meyer and Liu, 2014). Therefore, consistent and reproducible processing of the chromatin profiling data is essential in deriving meaningful information from the experimental data. Moreover, experiments can fail due to technical complexities. Comprehensive quality control will help to identify failed samples, and robust processing can facilitate reproducible analysis.

We developed CHromatin enrIchment ProcesSor (CHIPS) to standardize processing and quality control evaluation for ATAC-seq, ChIP-seq, and DNase-seq data following best practices (Bailey *et al*., 2013). CHIPS is a scalable and reproducible pipeline written in Snakemake (Köster and Rahmann, 2012). Encapsulated in a Conda environment, it can be executed in the local computing cluster engine or in the cloud computing settings such as Amazon AWS and Google Cloud. CHIPS has been used to analyze >1500 samples since 2016 within Dana-Farber Cancer Institute, and now serves as the standard processing pipeline for tumor ATAC-seq data from the Cancer immune Monitoring and Analysis Centers and Cancer Immunologic Data Commons (CIMAC-CIDC) trials (Chen *et al*., 2021).

## 2 Methods

### 2.1 Alignment and basic quality control

CHIPS takes FASTQ or BAM files as input and supports both single-end and paired-end data. To save time and resources, CHIPS subsamples 100,000 reads and uses them in the FASTQC module for basic quality control analysis. For aligning reads to the reference genome, FASTQ files are trimmed to remove adaptors and low-quality sequences using fastp (Chen *et al*., 2018) and then aligned by BWA-MEM (Li and Durbin, 2009) to generate sorted and deduplicated BAM files. After alignment, the mapping statistics, including the number of mapped and uniquely mapped reads, are then reported.

CHIPS carries out other basic quality control. The contamination profile reports the percentage of 100,000 reads that map to a contamination panel’s reference genomes. The contamination panel, specified by the user in a configuration file, includes dm3, Saccharomyces cerevisiae, E. coli, and mycoplasma of different types in addition to hg38, hg19, mm10, and mm9 genomes. We provide static reference files along with the installation of CHIPS. Users may add new assemblies to the contamination panel by adding the BWA index files. In addition, 4,000,000 reads are downsampled for calculating the PCR bottleneck coefficient (PBC). The PBC is the number of locations with exactly one uniquely mapped read divided by the number of uniquely mapped genomic locations. PBC ranges from 0-1, and a higher number indicates higher library complexity.

Particularly useful in the setting of ATAC-seq experiments, CHIPS also provides a fragment lengths distribution plot. ATAC-seq data with high quality should have fragment length peaks at < 100 bp nucleosome-free regions and show periodical enrichment at the 1- and 2-nucleosome lengths.

### 2.2 Peak calling and peak characteristics for quality control

Peaks represent regions of the genome that are enriched with aligned reads. The MACS2 (Zhang *et al*., 2008) algorithm is used to call peaks from uniquely sorted bam files. The minimum false discovery rate (FDR) cutoff for defining peak confidence is set to 0.01 by default but can be changed in the config.yaml file. A summary of the number of peaks, including those with a > 10 or > 20-fold increase relative to the background, is also reported describing the data quality. More peaks and a higher fraction of >10X peaks tend to indicate higher quality. Moreover, a read per million (RPM) normalized BedGraph signal track file generated by MACS2 is further converted to a BigWig file for visualization in the genome browsers more efficiently. A qualitative assessment of peak quality can be determined by static genome browser track views in the CHIPS output.

After peak calling, the fraction of reads in peaks (FRIP) scores is calculated to assess the samples’ quality. The FRIP score is the fraction of 4,000,000 subsampled reads that fall within the peak regions. FRIP score increases with sequencing depth, so a subsample of reads is used. The FRIP score indicates data’s signal-to-noise ratio, and a higher FRIP score indicates higher quality.

Certain characteristics of the peaks can be used to describe further the quality of the data. Peaks from a high-quality sample should have a high percentage of overlap with the known DNaseI sites. CHIPS overlaps the peaks with the union of the public DNaseI hypersensitive sites to determine the data’s quality. Moreover, high-quality peaks tend to be evolutionarily conserved across species. CHIPS plots the conservation plot across all peaks. The conservation plots of transcription factors typically show a high focal point around the peak summits, while histone modifications show bimodal peaks with a dip in the center.

### 2.3 Downstream analysis

Peak annotation is performed to describe how the peaks distribute across the genome. Specifically, CHIPS determines the proportions of peaks that overlap with promoters, exons, introns, or intergenic regions. Motif identification is carried out using Homer (Heinz *et al*., 2010). The top 5,000 most significant peak summits (ranked by the MACS P-value) are used for motif analysis. Finally, to determine which genes may be regulated by the peaks, a regulatory potential score is calculated for each gene using an exponential decay model implemented in LISA (Qin *et* al., 2020). LISA calculates regulatory potential scores that represent the cumulative influence of nearby peaks associated with each gene.

### 2.4 Output

CHIPS provides results files in txt and png forms inside well-structured folders and a dynamic HTML report summarizing quality control metrics at the sample level. An example report for TCGA-LUAD ATAC-seq data is available at http://cistrome.org/∼lentaing/chips/report/report.html. Documentation accompanying the CHIPS software describes the installation process and the structure of the analysis results and report directories. Due to the modular nature of the Snakemake workflow, the report can be customized to meet individual needs and easily expanded if new metrics are added. Furthermore, the same metrics are reported in the CistromeDB (Zheng *et al*., 2019) which facilitates comparisons of results with that resource.

## 3 Conclusion

Taken together, CHIPS performs quality control and reproducible processing of the chromatin profiling data generated from ATAC-seq, ChIP-seq, and DNase-seq experiments. CHIPS does not explicitly label samples as being “low” or “high” quality overall. We rely on the users to interpret information from multiple quality control features to determine which samples to include for further downstream analyses. CHIPS also does not provide downstream analyses comparing cases and controls. Downstream analyses depend on the biological context of the experiments and may consist of differential binding, motif analysis, and pathway analysis in the setting of chromatin profiling experiments. An independent Snakemake pipeline COBRA(Qiu *et al*., 2020) is designed for this purpose.

**Fig. 1.**
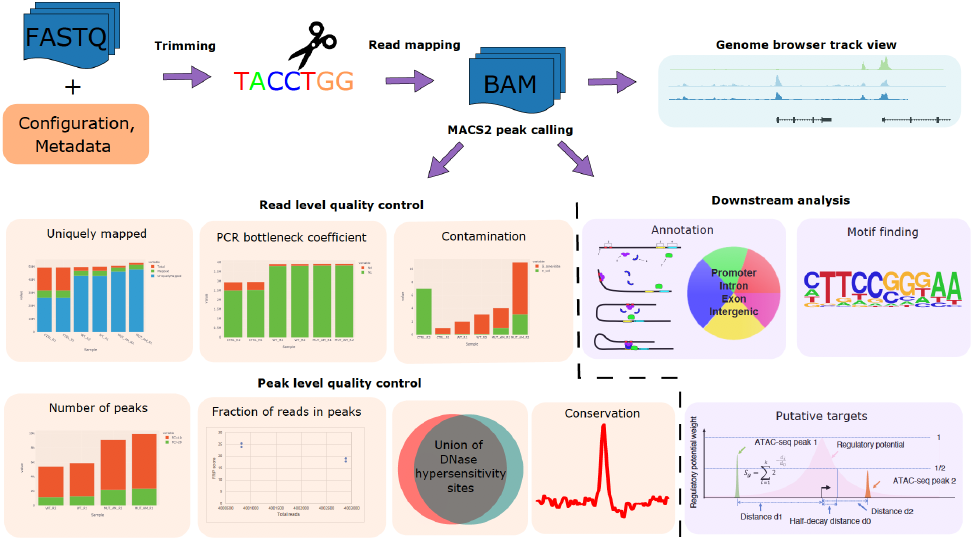
CHIPS workflow. The CHIPS pipeline is designed to perform robust quality control and reproducible processing of chromatin profiling data derived from ChIP-seq, ATAC-seq, and DNase-seq. The CHIPS pipeline includes basic steps of read trimming, read alignment, and peak calling. For quality control, it calculates metrics such as contamination profile, mapping statistics, the fraction of reads in peaks (FRIP) score, PCR bottleneck coefficient (PBC), overlap with union DNaseI hypersensitive sites (DHS), and peak evolutionary conservation. For downstream analysis, CHIPS carries out peak annotation, motif finding, and putative target prediction. The inputs to the pipeline are FASTQ/BAM format DNA sequence read files.

## Acknowledgements

We thank the Center for Functional Cancer Epigenetics and Molecular Biology Core Facilities at Dana-Farber Cancer Institute for valuable feedback on CHIPS.

